# Restriction-site-based enrichment coupled to adaptive sampling enables long-read transposon-insertion sequencing

**DOI:** 10.64898/2026.06.23.733934

**Authors:** Christian Jonas Lapp, Janek Weiler, Johannes Gescher

## Abstract

Long-read Transposon insertion sequencing is essential for linking genotypes to phenotypes, as short-read approaches struggle with complex genomic regions. Long-read sequencing demands high sequencing depth or targeted sequencing, which either requires extensive sample preparation or suffer from low net efficiency. We developed a simple, low-cost workflow combining enzymatic cleavage with adaptive sampling to drastically improve target recovery and simplify analysis. I-SceI restriction site was introduced into a mini-Tn5 transposon to generate a mutant library in *Cupriavidus necator*. Prior to Nanopore sequencing, gDNA was digested to introduce double-strand breaks precisely at insertion sites, followed by sequencing with adaptive sampling. The workflow was validated via a biofilm selection screen. Combining I-SceI digestion with adaptive sampling yielded an effective 11.3-fold enrichment (after normalizing for pore occupancy). High-depth sequencing covered 97.1% of all genes. Because reads consistently began at the exact insertion site, bioinformatic trimming was bypassed. The biofilm screen successfully identified enriched integration loci, uniquely mapped insertions within repetitive rRNA operons, and revealed co-selected, spontaneous secondary mutations.This method adds less than one hour to library preparation, reduces consumable costs by over 90%, and establishes a budget-friendly, simple and high-resolution long-read Tn-seq pipeline ideal for complex and multipartite genomes.

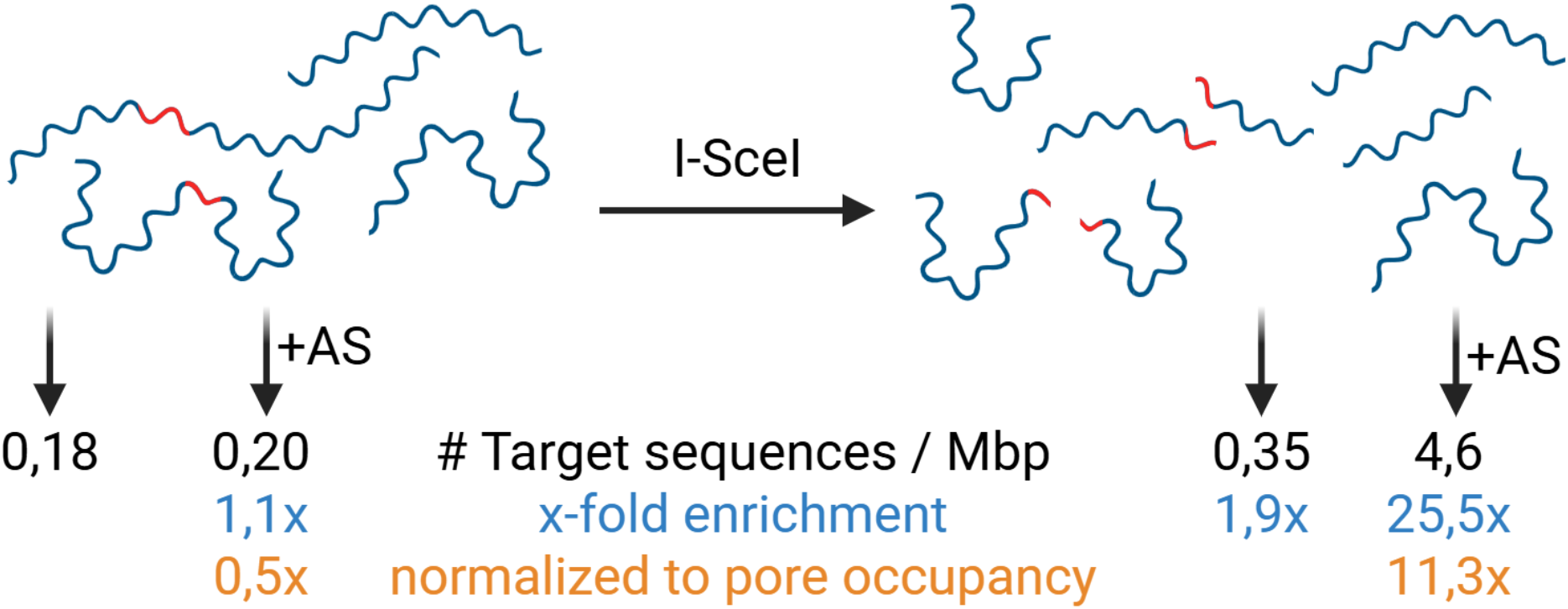

## Introduction

Linking genotypes to phenotypes has remained a central challenge in biology since the first complete genome sequences became available (1, 2). Systematic gene disruption approaches, such as targeted knockouts, have since been established to enable functional examination of genomes, yet phenotypic effects are often condition-dependent and require extensive experimental screening across diverse environmental contexts (3). Transposon mutagenesis has substantially accelerated functional genomics by enabling the generation of dense mutant libraries; however, identifying and quantifying insertion sites by sequencing at scale remains technically demanding. As a result, targeted transposon insertion sequencing (TIS) and related methods have emerged as powerful tools to associate gene disruptions with fitness phenotypes in a high-throughput manner (4, 5). These approaches reduce time and resource requirements by coupling pooled mutant libraries with next-generation sequencing, where each read corresponds to an insertion event. Classical Tn-seq protocols rely on restriction enzyme digestion near the transposon insertion site, followed by adapter ligation and PCR amplification, generating fragments well suited for short-read sequencing platforms (6). While efficient, these approaches are limited by short read lengths, which complicate mapping in repetitive or structurally complex genomic regions, and by PCR-induced biases that affect quantitative interpretation (7, 8).

Advances in sequencing technologies, particularly third-generation long-read platforms such as Oxford Nanopore and PacBio, offer new opportunities to overcome these limitations (9). Long reads enable improved resolution of repetitive regions, direct detection of structural variants, and, in some cases, identification of epigenetic modifications (10). However, applying long-read sequencing to transposon insertion mapping introduces new challenges, particularly the requirement for sufficient sequencing depth. Because each insertion event must be sampled individually, sequencing demands scale with both genome size and insertion library complexity, motivating the development of targeted enrichment strategies(10).

TIS approaches aim to increase the proportion of reads originating from the regions of interest. Amplicon sequencing is a robust and widely used strategy that relies on PCR amplification with primers specific to known sequences and is also compatible to long read seuencing. Although highly sensitive and effective for low-input samples, it is constrained by the requirement for known flanking regions, limited amplicon length, and amplification bias (10, 11). Hybrid capture methods address some of these limitations by enriching target sequences via probe hybridization, often achieving enrichment factors of 10^4^–10^5^ (12, 13). However, the typically low post-capture DNA yield necessitates amplification, reintroducing biases similar to those observed in amplicon-based approaches (14).

CRISPR/Cas9-based enrichment has emerged as a programmable alternative, enabling targeted cleavage of DNA sequences followed by adapter ligation (15, 16). This method avoids dependence on flanking sequences and supports long fragments, but practical limitations remain, including high DNA input requirements, challenges in optimizing guide RNA design, and potential interference of Cas9–DNA complexes with downstream sequencing processes. In nanopore sequencing, persistent Cas9 binding has been associated with reduced pore performance and throughput, which is why this methods support by oxford nanopore was eventually discontinued (16–18).

Adaptive sampling, a software-driven approach unique to Oxford Nanopore sequencing, enables real-time enrichment by selectively rejecting non-target molecules during sequencing (19, 20). This approach is highly flexible and does not require additional wet-lab enrichment steps. However, enrichment efficiency depends on rapid target recognition, which is influenced by fragment length and the position of the target sequence relative to the sequencing start site. In practice, enrichment factors vary widely, and the method can suffer from reduced sequencing yield due to pore blocking and repeated rejection events (21–23). Although combinations of adaptive sampling with upstream enrichment strategies, such as Cas9-mediated cleavage, have been proposed, practical implementations remain limited as no substantial enrichment of target reads could be obtained compared to nCATS alone and further developments of this method are regarded necessary (24).

In this study, we describe a simple and robust method, introducing just one additional step to the sample preparation with an additional lead time of less than one hour including hands-on time of 10 – 15 minutes. By introducing an additional restriction site into the transposon sequence, we established a workflow similar to Tn-seq that improves target enrichment in by >10x when combined with adaptive sampling. While, at the same time, simplifying the analysis workflow with a DNA input demand on par with standard sequencing protocols for budget-friendly Oxford Nanopore sequencing.

## Material and Methods

### Preparation of the transposon library

The transposon library was generated using the mini-Tn5 delivery plasmid pBAMD1-2, which was kindly provided by Víctor de Lorenzo and obtained from Addgene (plasmid #61564; RRID:Addgene_61564). The plasmid was delivered in *Escherichia coli* and purified using the Wizard® Plus SV Minipreps DNA Purification System (Promega, Mannheim, Germany). pBAMD1-2 is a standardized mini-Tn5 vector designed for broad-host-range transposon delivery in Gram-negative bacteria, making it suitable for insertional mutagenesis experiments.

Because the aim of this study was to generate stable chromosomal mutations without introducing additional functional elements, the original kanamycin resistance cassette of the transposon was retained, and only an additional I-SceI recognition site was introduced. The I-SceI site was inserted by inverse PCR using primers #4555 (GCGGCCGCGCGAATTCG) and #4556 (ATTACCCTGTTATCCCTAGGCCTAGGCGGCCTGAGA), with primer #4556 carrying the I-SceI recognition sequence as an overlap.

The inverse PCR was performed at 69 °C with 3% DMSO using a high-fidelity DNA polymerase (PCR-Biosystems Ltd, London, UK), and the product was purified using the Wizard® SV Gel and PCR Clean-Up System (Promega, Mannheim, Germany). The PCR product was then phosphorylated with T4 polynucleotide kinase (New England Biolabs, Frankfurt, Germany) at 37 °C for 20 min and ligated with T4 DNA ligase (New England Biolabs, Frankfurt, Germany) at room temperature overnight, according to the manufacturer’s instructions. The resulting plasmid was verified by gel electrophoresis, repurified using the Wizard® SV Gel and PCR Clean-Up System, and quantified using a NanoDrop 2000 spectrophotometer (Thermo Scientific, Waltham, USA).

The verified plasmid was introduced into the DAP-auxotrophic *E. coli* donor strain WM3064, which is commonly used for conjugative transfer of mobilizable plasmids because its DAP dependence allows counterselection after mating. Sequence-confirmed clones were obtained by Sanger sequencing (Eurofins Genomics, Ebersberg, Germany) using the Mix2Seq kit and compared to the *in-silico* plasmid sequence in Benchling (San Francisco, USA). After identification of a clone carrying the correct construct, the strain was used as the intermediate donor for conjugation into the *Cupriavidus necator* wild-type strain DSM 428 (H16).

To estimate the number of transformants required to cover insertions in non-essential genes, we assumed random insertion and calculated the library size needed to achieve at least 90% coverage of genes with an average length of 1 kb in a 7.4 Mb genome, following standard transposon-library design principles. Under these assumptions, at least 17,038 independent insertion events were required. Colony-forming units were determined by serial dilution of both strains at OD_600_ = 1.0 and plating on LB agar. For mating, both cultures were grown separately to OD_600_ = 0.6, and at least 1,000 colony-forming units of each strain were combined on 20 LB agar plates supplemented with 0.3 mM DAP and incubated for 2 days at 30 °C. This procedure minimized cell–cell contacts during conjugation. After incubation, cells were recovered by washing the plates with 20 mL fresh LB medium and replated onto 20 new plates containing 250 µg/mL kanamycin for selection. Colony numbers were counted after incubation, and 26,185 colonies were obtained, exceeding the required library size. The recovered library was then resuspended in 60 mL LB medium containing 250 µg/mL kanamycin and stored at −80 °C for downstream sequencing and phenotypic screening.

### Proof-of-concept experiment

The *C. necator* transposon library was cultivated as a biofilm to identify insertions that conferred a fitness advantage under these conditions. Biofilm growth was performed in a well-established microfluidic flow-cell system based on the setup described by Hansen et al. (2019). The experiment was conducted in four flow cells over 168 h until mature biofilms had formed. To avoid pre-selection before the experiment, no pre-culture was used for inoculation; instead, cells were taken directly from cryotubes and adjusted to an OD_600_ of 0.2. Inoculation was carried out over 2 h at a medium flow rate of 3 mL/h and an inoculate flow rate of 1 mL/h. Thereafter, cultivation was continued at a total flow rate of 4 mL/h and 30 °C throughout the experiment. After 168 h, the biomass was harvested by flushing the flow cells and stored at −80 °C. Genomic DNA from the biofilm populations was then isolated and re-sequenced for comparison with the original library.

### DNA isolation and preparation for sequencing

Genomic DNA was extracted using the DNeasy PowerBiofilm kit (Qiagen) according to the manufacturer’s instructions. Samples from the input library were obtained by washing approximately 26,000 colonies from LB agar plates into LB medium and collecting the suspension by centrifugation. Four biofilm samples were harvested from the microfluidic chip system after the growth period by flushing the chip with pre-warmed MBA solution from the DNeasy PowerBiofilm kit, and these samples were processed in the same way as the input library.

DNA quantity was determined using the Qubit dsDNA HS Assay Kit, and quality was assessed by NanoDrop 2000 absorbance measurements. For enzymatic enrichment, 4 µg of each DNA sample was digested with I-SceI (New England Biolabs) in CutSmart buffer for 4 h. Digested samples were cleaned using AMPure XP beads by adding an equal volume of beads, incubating for 10 min with rotation, pelleting on a magnetic rack, washing twice with 80% ethanol, and eluting in 12 µL ddH2O. Both digested and undigested samples were used for library preparation. For barcoding, each sample was individually labeled using the Native Barcoding Kit 24 V14 (Oxford Nanopore Technologies), and 600 ng in total were pooled for final library preparation according to the manufacturer’s protocol.

### Sequencing and data analysis

The library was loaded onto a MinION Mk1B and sequenced twice sequentially on the same R10.4.1 flow cell. In the second run, adaptive sampling was enabled to enrich reads matching the modified transposon sequence. A 350 bp FASTA target file containing flanking regions of the restriction site was used as the enrichment reference. Super-accurate basecalling was performed with MinKNOW release 25.05.14 and model v4.3.0, with adapter and barcode trimming enabled. Sequencing and basecalling were performed on a windows 11 workstation with 64 GB of RAM, Core i7-14700 and an NVIDIA Geforce RTX 4600 with 8 GB of RAM. Adaptive sampling is a nanopore-specific method for real-time enrichment of target sequences and has been shown to improve target representation in several applications, although performance depends strongly on target accessibility and fragment structure.

Data analysis was performed mainly in CLC Genomics Workbench 20.0.4 (Qiagen). Reads were identified as on target if they mapped to both, the transposon insertions sequence, and the genome. For further quantitative analysis of I-SceI-cut DNA, reads were trimmed against the modified Tn5 sequence using two adapter definitions representing the flanking regions of the I-SceI restriction site. Untrimmed reads were excluded. Trimmed reads were mapped with minimap2 to the *C. necator* reference genome (ASM479872v1) and alignment start positions evaluated for Transposon insertion positions. Comparative analysis of insertion events between samples was performed in Microsoft Excel and compared to genomic regions of annotated CDS.

Targeted-sequencing enrichment was calculated from the total sequencing depth and the number of reads that were successfully mapped to the modified Tn5 sequence and mapped to the reference genome. Because adaptive sampling reduces pore occupancy by rejecting off-target reads, an additional normalization was performed based on the number of available pores over time. For this normalization, occupied pores were related to the number of total healthy pores and compared with reads longer than 1,500 bp. A Python script was used to calculate pore occupancy and pore health over time; the script is provided in the supplementary material. Final values were combined, calculated, and validated in Microsoft Excel.

An additional sequencing run of the input Tn5 library was performed on a PromethION P2 Solo to obtain greater sequencing depth and to confirm genome-wide integration of the transposon after I-SceI digestion and adaptive sampling.

## Results

### 1. Targeted sequencing performance after I-SceI digestion and adaptive sampling

DNA from the transposon library was prepared for sequencing either directly or after I-SceI digestion, and each sample was sequenced on a MinION flow cell with adaptive sampling turned off and on. This yielded four sequencing datasets that were compared with respect to total sequencing output, the number of reads containing transposon-derived sequence, and pore occupancy during sequencing. The quantitative results are summarized in Table 1 and Figure 1.

**Table 1.**
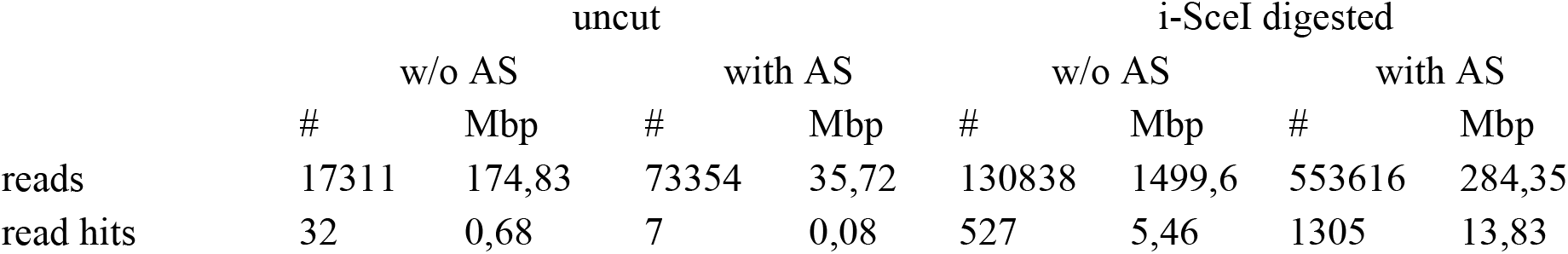
Comparison of sequencing results of different approaches for targeted sequencing. A standard sequencing approach without enzymatic digestion and without adaptive sampling is compared to adaptive sampling and to a pre-digestion with I-SceI. The number of reads and sequencing depth, are listed for raw sequencing data and for on-target reads.

|  | uncut |  |  |  | i-SceI digested |  |  |  |
| --- | --- | --- | --- | --- | --- | --- | --- | --- |
|  | w/o AS |  | with AS |  | w/o AS |  | with AS |  |
|  | # | Mbp | # | Mbp | # | Mbp | # | Mbp |
| reads | 17311 | 174,83 | 73354 | 35,72 | 130838 | 1499,6 | 553616 | 284,35 |
| read hits | 32 | 0,68 | 7 | 0,08 | 527 | 5,46 | 1305 | 13,83 |

**Figure 1.**
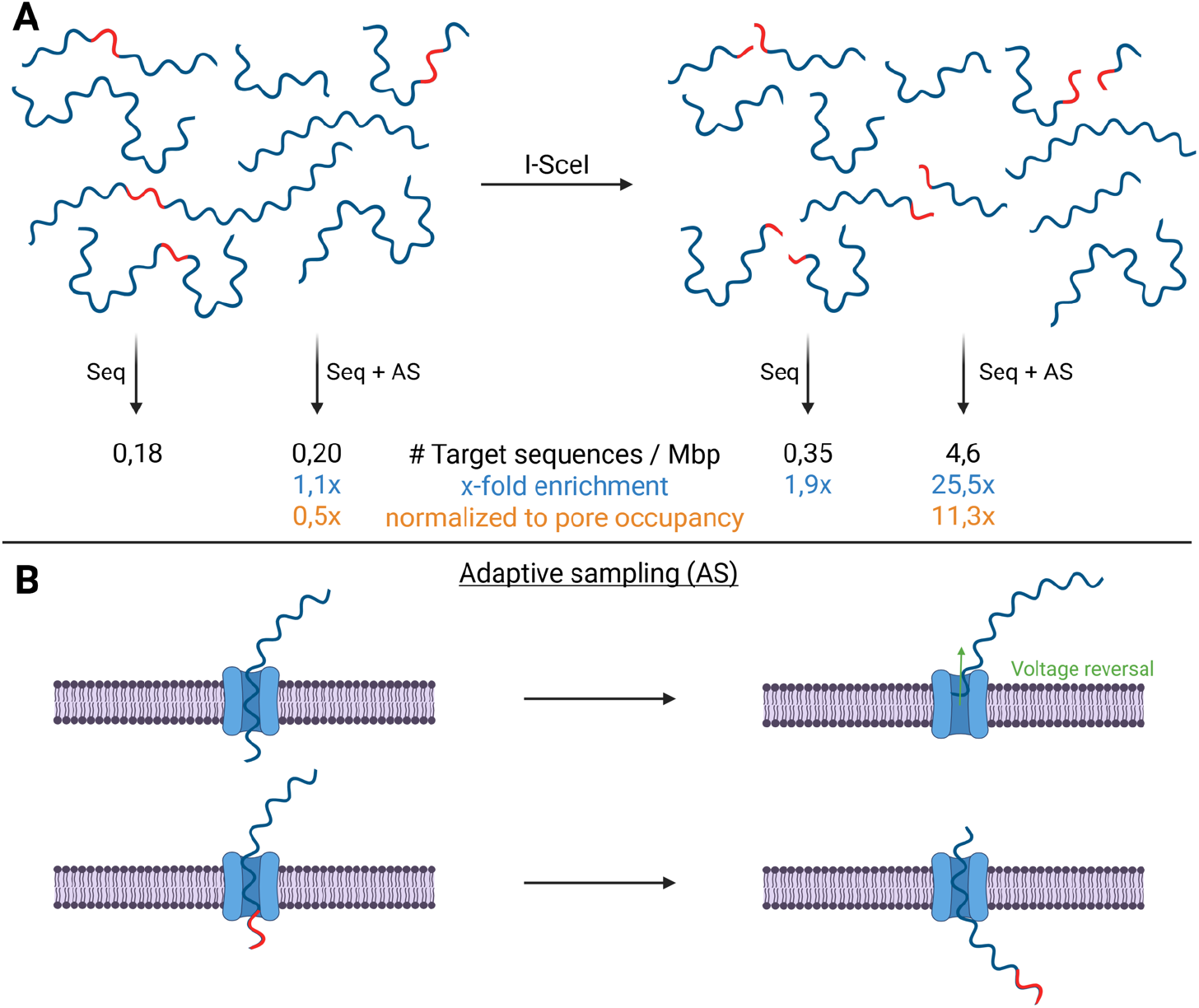
A: Comparison of targeted Sequencing using I-SceI digestion and adaptive sampling. First, native DNA is sequenced with and without adaptive sampling. In a second approach, cuts are introduced using an I-SceI restriction enzyme, and the DNA is sequenced in the same manner. The number of target sequences per Mbp are calculated and the comparative enrichment factor based on these numbers. For sequencing with AS an additional normalization was calculated based on the reduced sequencing depth which occurred due to adaptive sampling. B: Scheme of adaptive sampling recognition with target regions at the end of DNA fragments. Identification of the transposon sequence allows for rapid target recognition within the first few hundred basepairs.

For targeted long-read sequencing, adaptive sampling alone only modestly increased the fraction of target reads by 1.1-fold. However, because adaptive sampling also reduced overall sequencing output by temporarily occupying pores with rejected molecules, we additionally normalized enrichment to pore availability, following the principle that real-time read rejection can lower net throughput despite improving target representation. After this correction, adaptive sampling alone did not improve effective enrichment and instead resulted in a 51% reduction relative to sequencing without adaptive sampling.

In contrast, I-SceI digestion markedly improved target recovery. Digestion alone increased the number of fragments carrying transposon-associated sequence by 1.94-fold relative to the untreated control. Most notably, the combination of I-SceI digestion and adaptive sampling produced the strongest enrichment, yielding approximately a 25.5-fold increase in target reads relative to native sequencing without enrichment. Even after normalization for pore loss during adaptive sampling, the combined workflow still produced an 11.3-fold effective enrichment of target sequence. These results indicate that enzymatic pre-fragmentation substantially improves the performance of adaptive sampling for transposon-targeted nanopore sequencing.

A schematic overview of the workflow and the corresponding enrichment outcomes is shown in Figure 1.

### 2. Genome-wide coverage of the transposon library

To test whether the optimized workflow was sufficient for library-scale insertion mapping, the transposon pool was sequenced on a PromethION flow cell in two independent samples and barcodes using I-SceI digestion combined with adaptive sampling. This run produced 38.45 Gbp of sequencing output and 145,894 target reads, corresponding to approximately one target read every 51 bp. At this depth, the experiment was theoretically sufficient to detect insertions across all non-essential genes in the genome.

After mapping to the *Cupriavidus necator* reference genome, insertion sites were assigned to annotated genes. Of 6,881 genes, 6,682 were hit at least once (97.1%), 6,591 at least twice (95.8%), and 6,311 at least three times (91.7%). Fewer than 1% of genes were hit 75 times or more often, indicating that the library coverage was broad but not uniformly distributed. The insertion landscape was visualized in a CIRCOS plot, and replicon-specific coverage patterns were analyzed separately to account for differences among the three replicons (Figure 2 and Figure 3).

**Figure 2.**
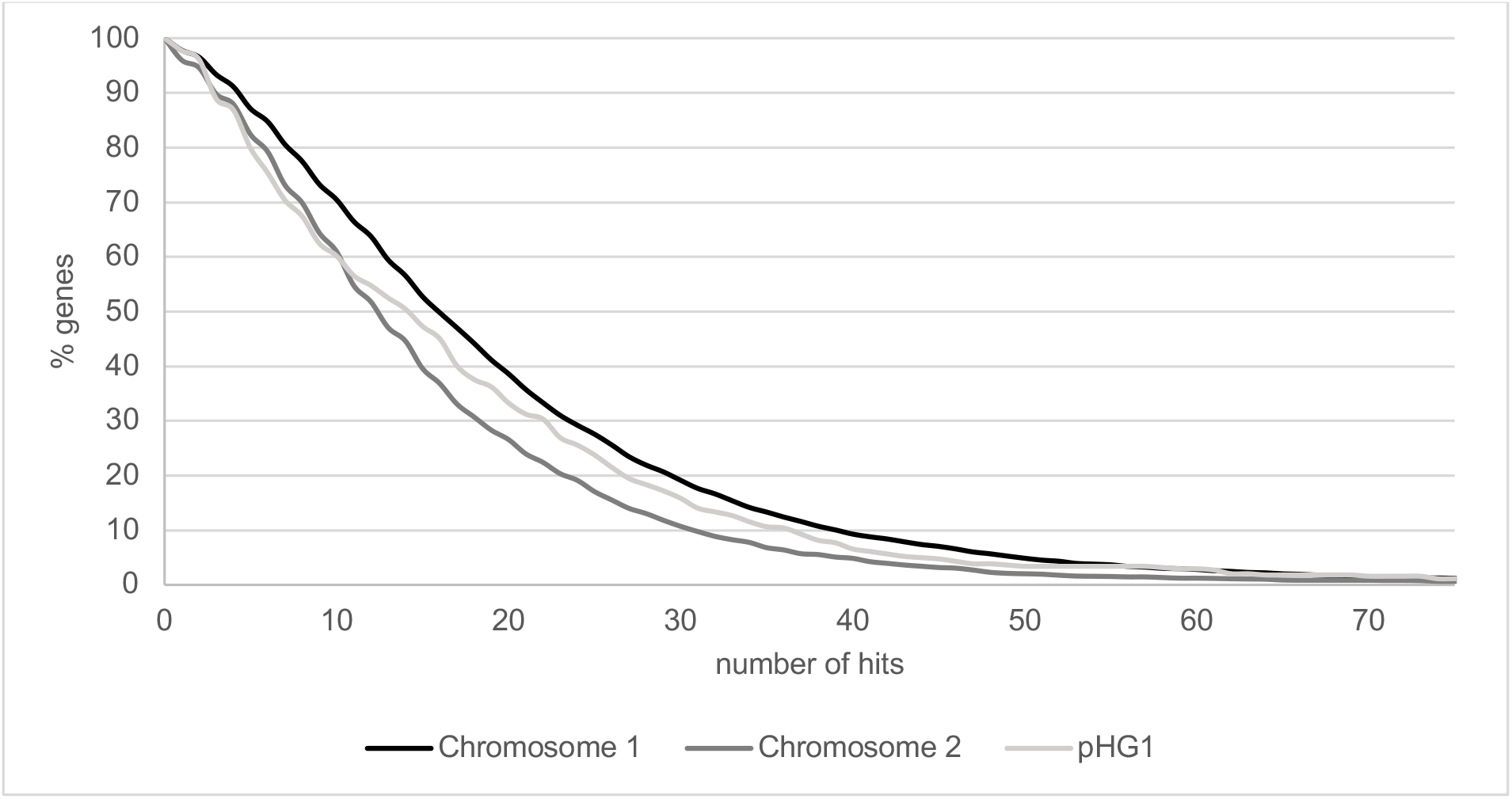
Number of insertion hits for genes on C. necator replicons. The proportion of genes are shown on the y-axis, the number of identified insertion hits on the x-axis. A separate presentation for the three replicons is shown to distinguish insertion frequencies.

**Figure 3.**
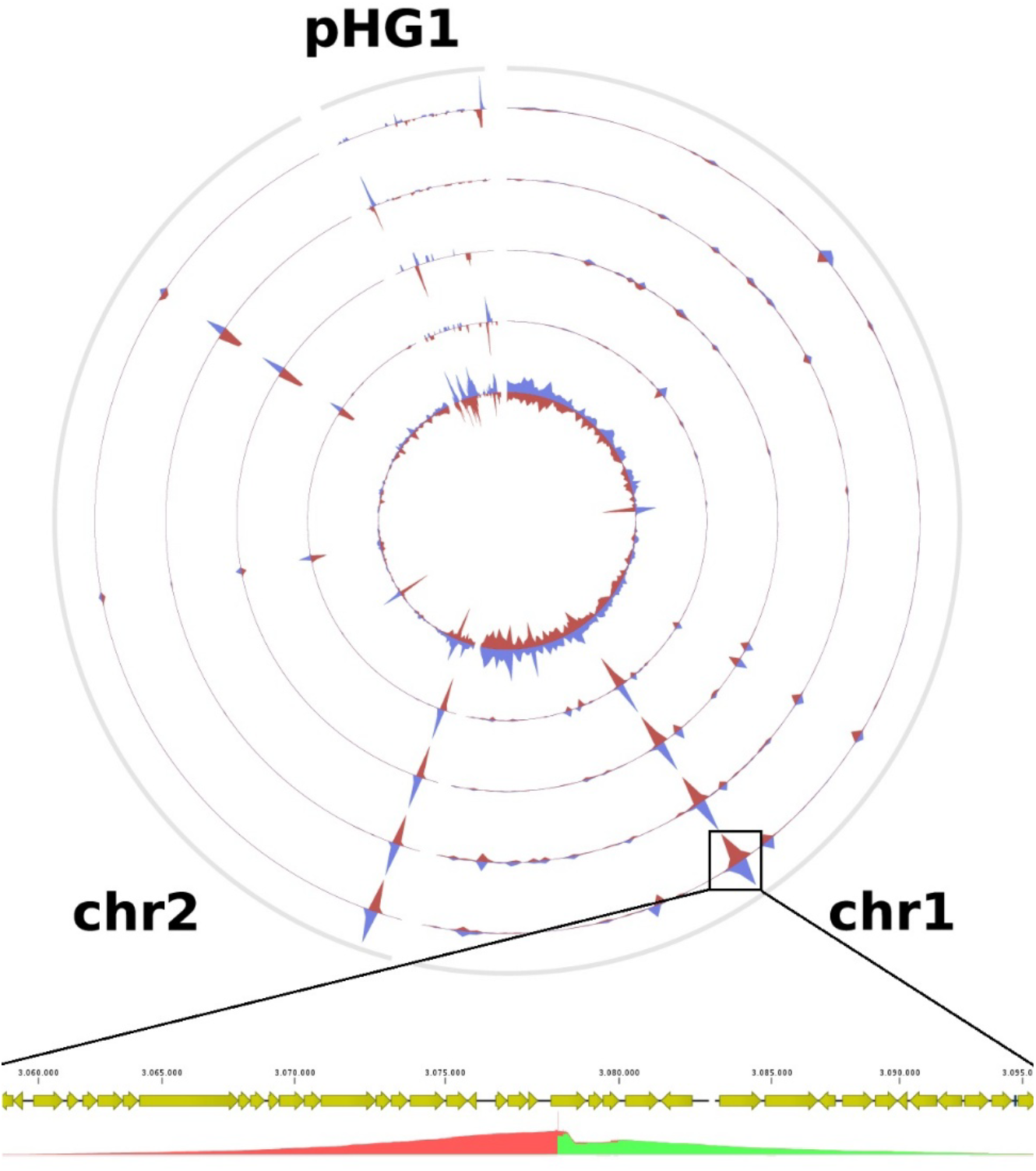
Circos plot that shows the mapping of target read mappings to the three replicons. The five circles represent mappings from three independent sequencing samples. The most inner circle derives from reads of the transposon library, while the outer four circles derive from a quadruplicate of biofilm cultivation experiments. A mapping result from the region of the gene ribBA from one of the biofilm replicates is shown on the bottom. Coverage by forward reads are colored in green, revers in red.

Insertion frequencies differed between the three replicons of *C. necator*. Chromosome 1 showed the highest average number of hits per gene, with 19.3 insertions per gene, followed by plasmid pHG1 with 17.0 hits per gene and chromosome 2 with 15.0 hits per gene. Despite these differences in insertion density, the proportion of genes hit at least once was highly similar across replicons, reaching 97.8% for chromosome 1, 96.0% for chromosome 2, and 97.7% for pHG1.

This pattern suggests that the library was broadly distributed across the genome but that local insertion frequencies were influenced by replicon-specific features.

### 4. Enrichment of biofilm-selected mutants

To identify transposon insertions associated with biofilm fitness, the transposon library was subjected to quadruplicate selection under biofilm-forming conditions and analyzed by MinION sequencing. Across the biofilm-selected samples, 12 genes were hit more than 100 times, 81 genes between 10 and 100 times, and 1,069 genes no more than 10 times. The resulting insertion profile is shown in Figure 3.

Compared with the input library, the biofilm-selected populations displayed a shifted insertion distribution, with more frequent targeting of chromosome 2. Although chromosome 1 had higher average insertion counts in the input library, chromosome 2 became enriched after biofilm selection. This shift was especially apparent among the 27 genes hit more than 50 times, of which only 4 were located on chromosome 1, whereas 23 were located on chromosome 2.

### 5. Read-level identification of insertion sites and local mutations

A benefit of this workflow is the insertion identification, where enzymatic digestion generated reads with alignment start positions at the precise transposon integration site as shown in Figure 3. The alignment pattern produced an interval of approximately 9 bp with double coverage, consistent with the expected target-site duplication generated by Tn5-mediated integration. A representative mapping example is shown in Figure 3.

Additionally, we wanted to investigate how reliable detection of integrations in repetitive genomic regions benefits from long-read sequencing and mapping. We searched for insertions in rRNA operons, where short-read approaches would often yield ambiguous placement. A rare insertion after selection to biofilm growth was detected in gene E6A55_RS17930, where nine reads supported an integration event. Two of these reads were only about 230 bp long and mapped ambiguously to multiple rRNA loci as would be expected from shot read sequencing. The remaining seven reads were several kilobases long and uniquely resolved the insertion site.

Another unexpected benefit of long-read mapping was discovered by simple variant detection analysis of the target reads mapping. These revealed two closely spaced mutations approximately 12 kb upstream of *ribBA* in three of four biofilm samples. These mutations were located in the genomic region highlighted by the mapping analysis in Figure 3 and were predicted to cause either a frameshift or a premature stop codon, thereby inactivating gene E6A55_RS14495 (pilY2). Notably, no transposon insertion was detected at this site in any of the experiments, suggesting that the observed sequence changes represent spontaneous secondary mutations which are linked to the corresponding transposon insertion in *ribBA*. Approximately 80% of reads spanning this genomic region carried the additional mutations, suggesting strong co-selection of a second loss-of-function event during biofilm growth.

## Discussion

Transposon insertion sequencing (TIS) is among the most widely used methods for conducting functional genetic screens in bacteria at a genome-wide scale (4). The random integration of transposons, followed by quantification of insertion sites through high-throughput sequencing, enables screening of nearly all possible beneficial gene disruptions in growth selection experiments with minimal effort. The key to detecting transposon insertions lies in targeted sequencing approaches that direct sequencing reads exclusively or predominantly toward insertion sites. While early methods such as Tn-seq and TraDIS (4, 6, 25) were designed for short-read sequencing technologies, emerging long-read technologies of the third generation (PacBio and ONT) overcome short-read-specific limitations, such as resolving insertions within repetitive genomic sequences (26).

Building on modern targeted sequencing strategies, several protocols for TIS on the nanopore sequencing platform have been reported. LoRTIS represents a modern PCR-based approach modified from TraDIS in which amplicon sequencing is used to generate long reads for TIS, offering the opportunity to resolve difficult genomic regions (26, 27). LoRTIS provides fragment lengths exceeding 1,200 bp with a yield of 4.2 to 7.6 million on-target reads from a single MinION run, which is approximately 100-fold more compared to our dataset. However, because this method is PCR-based, results are not directly comparable: amplicons represent copies of the same fragments, and PCR bias arising from primer annealing efficiency, adapter ligation efficiency, and fragment length affects the actual distribution of detected insertion loci. For better comparison to assess the sequencing efficiency is the genome-wide coverage. An indirect indicator is the number of genes with no detected insertions, which largely reflects putative essential genes. For LoRTIS, 498 genes (11%) showed no insertions in an *E. coli* library, of which 311 were verified by orthogonal sequencing methods (26). In our dataset, 199 genes (2.9%) showed no insertion, which is consistent with previously published short-read data (28, 29). Even though statistical bias can partially be circumvented in PCR-based system by usage of UMIs (30), a high number of on-target hits is not sufficient to ensure complete genome coverage, and is therefore not directly comparable to native sequencing methods.

To our knowledge, no successful PCR-free hybrid capture methods for targeted sequencing has been reported, leaving Cas9-based methods the only option for targeted, unbiased native DNA sequencing. Since its development nCATS has been widely applied (31, 32) and reported to achieve 0.5–15% on-target reads (16) and up to 724-fold (33) enrichment in eukaryotic cell lines. An adaptation of this method specifically developed for TIS in bacteria is nCATRAS, which was designed to address amplification bias and is the most comparable approach to the method we present here. The authors of nCATRAS noted challenges in optimizing the library preparation but ultimately developed a protocol that achieved significant sequence enrichment with 23% on-target reads. This corresponds to 460,000 on-target reads obtained from three MinION flow cells, which typically yield 60–100 Gbp (8) and approximately equals the output of a PromethION flow cell, from which we yielded 146,000 on-target reads. nCATRAS represents a highly optimized implementation of nCATS and demonstrates the differences that arise when comparing target sequencing approaches in eukaryotic and prokaryotic samples in different context. In most nCATS approaches a region of defined sequence length is targeted where bordering cuts are introduced and the coverage of this area is observed. Methods with single-end cuts show lower enrichment rates (31). As insertion sites are investigated in TIS coverage is not applicable which is why the number of on-target reads is most relevant. The number of on-target reads also highly depends on the genomic background which is far larger in eukaryotic genomes where a great variance of 100 to 10,000 on-target reads from one MinION flow cell can be observed (16, 33).

On-target reads in our dataset were assessed using the same criteria as in nCATRAS: reads were filtered by a minimum length of 1,500 bp and required to map to both the transposon sequence and the genome. On this basis, nCATRAS achieves approximately a 3-fold higher enrichment rate. That said, Cas9-based sequencing methods require labor-intensive sample preparation, including guide RNA design and synthesis, large DNA input quantities, size selection, and carry a substantial risk of sequencing output reduction due to pore clogging. The authors themselves concluded that PCR-based TIS remains preferable when weighing the effort against the results (8). Given that Cas9-based approaches were developed for earlier generations of nanopore sequencing chemistry (R9.4), and limitations of these approaches ultimately led Oxford Nanopore Technologies to discontinue support for this technique in current chemistry generations (R10.4), but the promise of native long reads for generating information-rich data persists, we chose to develop a comparable method requiring substantially less sample preparation and avoiding the pore-clogging issues associated with Cas9 complexes.

This was achieved by combining a well-established and specific restriction enzyme with software-driven enrichment adaptive sampling which increases methodological robustness, reduces consumable costs and further reduces preparation time to less than an additional hour. Although adaptive sampling is a powerful enrichment tool and holds particular promise for TIS, expectations for this technology have declined considerably since its introduction, as its effectiveness is highly context-dependent and reported enrichment rates vary widely across studies (21–23). In this work, adaptive sampling alone resulted in a substantial loss of sequencing efficiency: we observed a low enrichment factor (1.1-fold) alongside a 56% reduction in total sequencing output. This is because the transposon insertion sequence is small relative to total read length, and sequencing decisions are made within the first few hundred base pairs, causing many reads that would ultimately contain an insertion site to be prematurely rejected. As the limitations of Cas9-based targeted sequencing persist — and are further exacerbated — when combined with adaptive sampling (24), this combination was deemed infeasible. However, the introduction of defined double-strand breaks into the DNA substantially simplifies downstream data analysis. Because sequencing reads consistently initiate at the insertion site, alignment can be performed directly on raw reads without prior trimming of transposon sequences; the transposon portion remains unmapped and is therefore excluded from alignment files. As a result, insertion sites can be identified efficiently by extracting read start positions from the alignment output generated during adaptive sampling. The same mapping data can further be used to visualize and validate insertions, including in complex genomic regions (Fig. 3), and to detect sequence variants co-occurring with insertion events.

We demonstrate that incorporation of an I-SceI restriction site alone, followed by enzymatic digestion resulted in a 1.94-fold enrichment of the target sequences. The additional application of adaptive sampling further increased enrichment to 11.3-fold. This combined approach substantially reduces the required sequencing depth and, consequently, consumable costs by more than 90% making it suitable for applications such as biofilm fitness experiment. Importantly, it mitigates a key limitation of adaptive sampling alone, namely delayed target recognition that allows off-target molecules to occupy sequencing pores. I-SceI digestion showed no detectable adverse effects on pore health. While adaptive sampling reduced sequencing output by an average of 56% in MinION experiments, a total yield of 38.4 Gbp was achieved on a PromethION flow cell, consistent with reported outputs of approximately 100+ Gbp under standard conditions and 37–52 Gbp with adaptive sampling enabled. When comparing the PromethION sequencing reads to MinION datasets we observed a slight enrichment decrease to a factor of 3.8 reads/Mbp instead of 4.6 reads/Mbp. This observation may be due to run-to-run variations, but an increased decision time in adaptive sampling could be correlated, as average read rejection occurred after 600 bp compared to 424 bp on MinION runs. This probably corresponds to the increased amount of 2675 channels in PromethION flow cells compared to the 512 in MinION flow cells and therefore an increased processing demand. Effectiveness may therefore partially be dependent on hardware configuration and workload of the sequencing computer. As our setup presents a rather average configuration, at least similar results can be expected from modern systems.

To highlight the benefits of native long reads in TIS we investigated several aspects, which are not visible with short-read data. A key outcome of this work is that long-read targeted sequencing enables high-resolution mapping of transposon insertions across the multipartite *C. necator* genome with multiple gene copies over its different replicons, where conventional Tn-seq approaches may collapse distinct insertion events into ambiguous mappings. Long-read sequencing therefore provides an additional level of resolution that is critical when the biological interpretation depends on precise genomic context rather than gene-level insertion frequencies alone. The ability to detect insertion sites within repetitive rRNA-associated regions demonstrates the strength of long-read sequencing in this workflow.

In addition to transposon insertion sites, we detected point mutations in a subset of sequencing reads, carrying at least one additional variant. These mutations were non-randomly distributed, occurring at elevated frequency near *ribBA*. Whether these variants arose as adaptive secondary mutations during biofilm selection, or represent pre-existing variation in the library, cannot be determined from the sequencing data alone. Regardless of their origin, their co-occurrence with insertion events illustrates a key advantage of native long-read sequencing over short-read TIS methods: the ability to detect multiple co-occurring genomic changes within a single read. Functional reconstruction of candidate mutations, individually and in combination, will be necessary to determine whether they contribute to the observed fitness phenotype. This is consistent with the expectation that strong selective pressures promote multi-locus adaptation, particularly in structured environments such as biofilms. Future work should focus on reconstructing these candidate mutations, individually and in combination, to determine whether they confer additive or synergistic fitness effects.

In addition, replicon-specific differences in insertion density demonstrate the complexity of transposon accessibility across the genome. Although nearly all genes were disrupted at least once, the three replicons of the genome exhibited distinct average insertion frequencies, indicating non-uniform insertion patterns. These differences likely arise from a combination of biological factors, such as gene essentiality, chromosomal organization, transcriptional activity, and DNA topology which were discussed before (34, 35). Accounting for these biases is important, as they can affect the inferred fitness landscape of mutant libraries and may otherwise confound data interpretation. The biofilm selection experiment illustrates the applicability of this approach for phenotype-driven enrichment. The observed shift in insertion frequency toward chromosome 2 following biofilm growth suggests that this replicon contains loci that are either more permissive to beneficial disruption or play a disproportionate role in biofilm-associated adaptation. However, these findings should be interpreted as an enrichment of candidate loci rather than definitive evidence of causality, as functional validation remains necessary.

These findings demonstrate the applicability and value of our method to even more complex, multipartite genomes. The transfer of this method to eukaryotic genomes would require corresponding adjustments, particularly regarding the integration of an I-SceI restriction site into eukaryotic transposon systems. The data from nCATS in human cell lines (16) and nCATRAS in *E. coli* (8) suggest that enrichment performance of Cas9-based targeted sequencing depends mostly on genome size, but is broadly comparable across these divergent organisms, providing an indirect reference point for expectations in more complex genomes. Some properties of our approach may even facilitate adaptation to larger genomes: the absence of guide RNA design requirements reduces organism-specific optimization, reduced off-target reactions of Cas9 (36), and the reliance on a defined enzymatic digestion step rather than Cas9 complex formation avoids the pore-clogging issues that are likely to be exacerbated by the higher DNA complexity of eukaryotic samples. However, whether these advantages translate into comparable enrichment in practice remains to be demonstrated, and we consider this an important direction for future development.

In conclusion, we present a method that is extremely easy to apply to general library preparation workflows, resulting in greater accessibility and reduced costs compared to most comparable Cas9-based approaches, even if the approximately 3-fold higher demand for sequencing depth is taken into account. Our findings demonstrate the advantage of native long reads over PCR-based and short reads in TIS in regards of sensitivity, specifity and statistical analysis, especially in context of multipartite genomes.

Finally, it is important to note that our method is particularly suited for transposon insertion sequencing, as it relies on the integration of a specific restriction site within the transposon sequence. In contrast, nCATS offers greater flexibility for targeting a broader range of sequences through the use of programmable nucleases. Overall, this study represents a step toward more efficient targeted sequencing strategies by demonstrating the advantages of combining defined DNA fragmentation with adaptive sampling, while also highlighting opportunities for expanding target versatility in future developments.

## Data Availability Statement

The raw sequencing data generated during this study have been deposited in the NCBI Sequence Read Archive (SRA) under accession number PRJNA1461446. The Python script used for analyzing pore occupancy and all other relevant data analysis pipelines are available in the supplementary material.

## Acknowledgements

The graphical abstract (Gescher, J. (2026) https://BioRender.com/kloaqn2) and Figure 1 (Gescher, J. (2026) https://BioRender.com/ockzcxz) were created in BioRender.

The authors used Claude (Anthropic, claude.ai, Sonnet 4.6) to assist with language editing and revision of the manuscript text. All content was critically reviewed, verified, and approved by the authors, who take full responsibility for the accuracy and integrity of the work

