## Supplements for "Restriction-site-based enrichment coupled to adaptive sampling enables long-read transposon-insertion sequencing"

Transposon sequences for trimming of raw reads:

ATCCCTAGGCCTAGGCGGCCTGAGACACAAAGATGTGTATAAGAGACAG

AACAGGGTAATGCGGCCGCGCGAATTCGAGCTCGGTACCCGGGGATCCTCTAGAGTCGA
CCTGCAGGCATGCAAGCTTGCGGCCGCCAAAGCCCGCCGAAAGGCGGGCTTTTCTGTATT
TAAATTTGTGTCTCAAAATCTCTGATGTTACATTGCACAAGATAAAAATATATCATCATGA
ACAATAAAACTGTCTGCTTACATAAACAGTAATACAAGGGGTGTTATGAGCCATATTCAG
CGTGAAACGAGCTGTAGCCGTCCGCGTCTGAACAGCAACATGGATGCGGATCTGTATGGC
TATAAATGGGCGCGTGATAACGTGGGTCAGAGCGGCGCGACCATTTATCGTCTGTATGGC
AAACCGGATGCGCCGGAACTGTTTCTGAAACATGGCAAAGGCAGCGTGGCGAACGATGT
GACCGATGAAATGGTGCGTCTGAACTGGCTGACCGAATTTATGCCGCTGCCGACCATTAA
ACATTTTATTCGCACCCCGGATGATGCGTGGCTGCTGACCACCGCGATTCCGGGCAAAAC
CGCGTTTCAGGTGCTGGAAGAATATCCGGATAGCGGCGAAAACATTGTGGATGCGCTGGC
CGTGTTTCTGCGTCGTCTGCATAGCATTCCGGTGTGCAACTGCCCGTTTAACAGCGATCGT
GTGTTTCGTCTGGCCCAGGCGCAGAGCCGTATGAACAACGGCCTGGTGGATGCGAGCGAT
TTTGATGATGAACGTAACGGCTGGCCGGTGGAACAGGTGTGGAAAGAAATGCATAAACT
GCTGCCGTTTAGCCCGGATAGCGTGGTGACCCACGGCGATTTTAGCCTGGATAACCTGAT
TTTCGATGAAGGCAAACTGATTGGCTGCATTGATGTGGGCCGTGTGGGCATTGCGGATCG
TTATCAGGATCTGGCCATTCTGTGGAACTGCCTGGGCGAATTTAGCCCGAGCCTGCAAAA
ACGTCTGTTTCAGAAATATGGCATTGATAATCCGGATATGAACAAACTGCAATTTCATCT
GATGCTGGATGAATTTTTCTAAGACCCTTGTCTAATTAATTGCGGACCCTAGAGGTCCCCT
TTTTTATTTTAAAAATTTTTTCACAAAACGGTTTACAAGCATAAAATCTCTGAAGATGTGT
ATAAGAGACAG

DNA sequences for adaptive sampling:

gcggccgcgcgaattcgagctcggtacccggggatcctctagagtcgacctgcaggcatgcaagcttgcggccgccaaagcccgccgaaaggc
gggcttttctgtatttaaatttgtgtctcaaaatctctgatgttacattgcacaagataaaaatatatcatcatgaacaataaaactgtctgcttacataaaca
gtaatacaaggggtgttatgagccatattcagcgtgaaacgagctgtagccgtccgcgtctgaacagcaacatggatgcggatctgtatggct

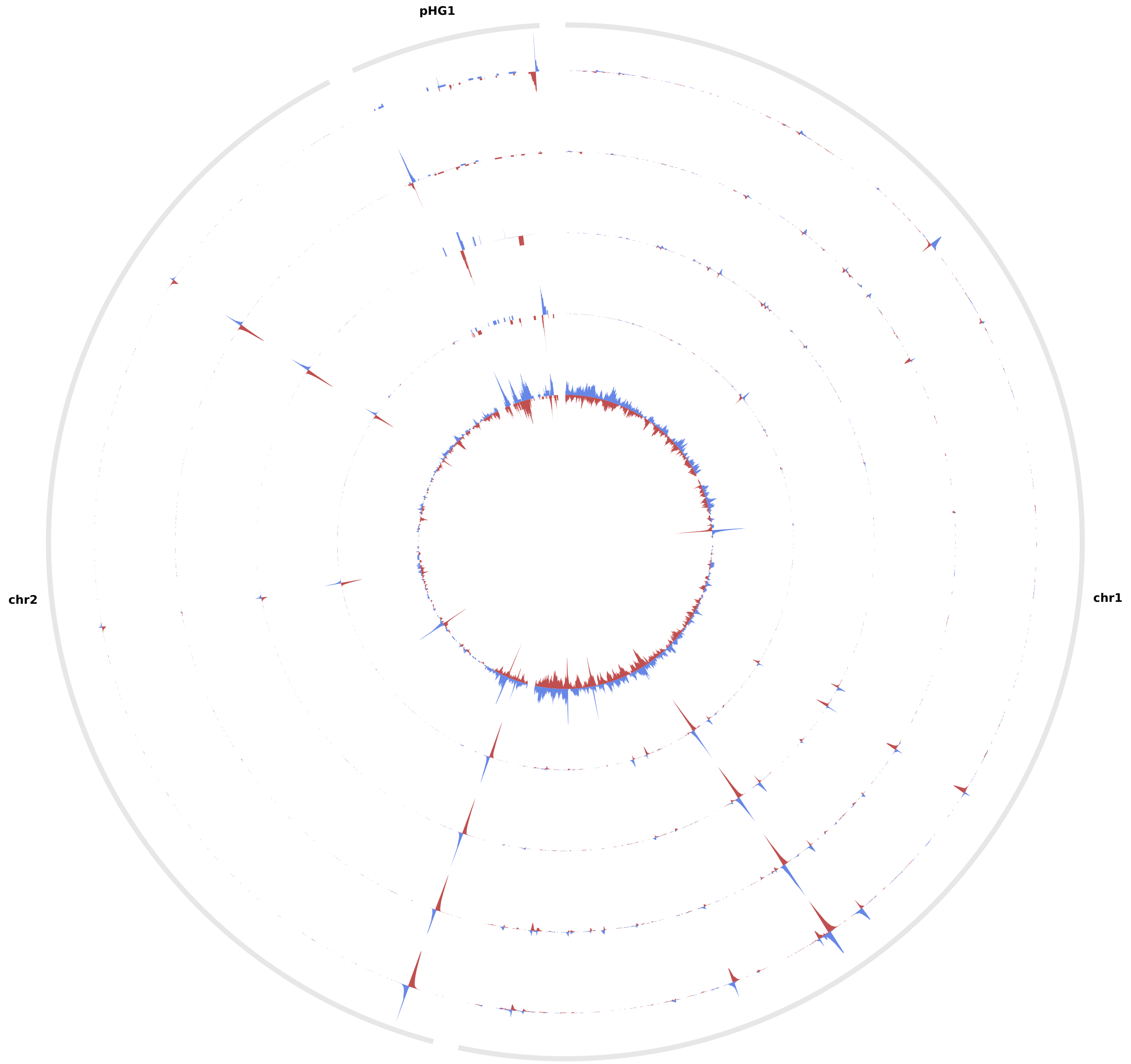


*Figure S1: Full scale svg circus plot of sequencing read mapping to the pangenome of Cupriavidus necator.*

Python script to calculate pore occupancy:

```
import pandas as pd
import numpy as np
import os

file_summary = "sequencing_summary_*.txt"
file_scan = "pore_scan_data_*.csv"

min_read_length = 1500
sequencing_speed = 400


scan_stats = pd.DataFrame(columns=['hour_bucket', 'Total_Good_Pores',
'Available_Channels'])
seq_stats = pd.DataFrame(columns=['hour_bucket', 'Active_Sequencing_Pores'])

if os.path.exists(file_scan):
    try:
        cols_scan = ['seconds_since_start_of_run', 'mux_scan_assessment', 'channel']

        df_scan = pd.read_csv(file_scan, usecols=cols_scan, dtype={'mux_scan_assessment': str,
'channel': int})

        mask_healthy = df_scan['mux_scan_assessment'].str.contains('single', case=False,
na=False)
        df_healthy = df_scan[mask_healthy].copy()

        df_healthy['hour_bucket'] = (df_healthy['seconds_since_start_of_run'] / 3600).astype(int)

        scan_stats = df_healthy.groupby('hour_bucket').agg(
            Total_Good_Pores=('mux_scan_assessment', 'count'),
            Available_Channels=('channel', 'nunique')
        ).reset_index()


if os.path.exists(file_summary):
    try:
        header = pd.read_csv(file_summary, sep='\t', nrows=0).columns.tolist()
        cols_needed = ['start_time', 'sequence_length_template']
        if 'duration' in header: cols_needed.append('duration')

        df_sum = pd.read_csv(file_summary, sep='\t', usecols=cols_needed)

        if 'duration' not in df_sum.columns:
            df_sum['duration'] = df_sum['sequence_length_template'] / sequencing_speed

        df_sum = df_sum[df_sum['sequence_length_template'] > min_read_length].copy()
        df_sum['hour_bucket'] = (df_sum['start_time'] / 3600).astype(int)

        seq_stats_raw = df_sum.groupby('hour_bucket')['duration'].sum().reset_index()
        seq_stats_raw['Active_Sequencing_Pores'] = (seq_stats_raw['duration'] / 3600).round(1)
```

```
        seq_stats = seq_stats_raw[['hour_bucket', 'Active_Sequencing_Pores']]


if scan_stats.empty and seq_stats.empty:
    print("no data")
else:
    final_df = pd.merge(scan_stats, seq_stats, on='hour_bucket',
how='outer').sort_values('hour_bucket').fillna(0)

    print("\n" + "="*100)
    print(f"{'Std':<4} | {'Pores (total)':<15} | {'Channels (Limit)':<15} | {'Active (Avg)':<12} |
{'saturation %':<20}")
    print("-" * 100)

    last_channels = 0

    for index, row in final_df.iterrows():
        h = int(row['hour_bucket'])
        pores = int(row['Total_Good_Pores'])
        channels = int(row['Available_Channels']
        active = row['Active_Sequencing_Pores']

        if channels > 0:
            last_channels = channels

        curr_limit = last_channels if channels == 0 else channels

        if curr_limit > 0:
            percent = (active / curr_limit) * 100

            marker = ""
            if percent > 85: marker = " 🔥 (Voll)"
            elif percent < 10: marker = " 📉 (Leer)"

            perc_str = f"{percent:.1f}% {marker}"
        else:
            perc_str = "-"

        if curr_limit == 0 and active == 0:
            continue

        print(f"{h:<4} | {pores:<15} | {curr_limit:<15} | {active:<12} | {perc_str:<20}")

    print("="*100)
    final_df.to_csv("pore_occupancy.csv", index=False)
```
